# Effects of metsulfuron-methyl on aquatic plant (*Lemna gibba* L.) and recovery from after prolonged exposure under rice cropping conditions

**DOI:** 10.1101/000638

**Authors:** Rathinasamy Raja Rajeswari, Atmakuru Ramesh

**Affiliations:** Department of Analytical Chemistry, International Institute of Bio-technology and Toxicology (IIBAT), Padappai Chennai, 601 301, Tamil Nadu, India

**Author notes:** Corresponding Author. /27174266; E-mail Addresses.

**Keywords:** Metsulfuron-methyl, *Lemna gibba*, Pigment content, Biomarker

## Abstract

The effects and potential recovery of aquatic plant *Lemna gibba* exposed to a sulfonyl urea herbicide metsulfuron-methyl (MSM) for 120 days under rice cropping condition was investigated. The frond number was decreased by day 15 at the concentration 11 µg/L and 100% inhibition on growth rate of *Lemna* was observed. Continuous decrease of frond number by day 50 at below the detectable level of residues exhibited symptoms (chlorosis) of MSM toxicity. Toxicity was assessed on the basis of toxicity index (TI) value, growth rate, yield and pigment contents (chlorophyll, carotene, total carotenoid and xanthophyll) of treated samples compared with untreated control. The observed value of 0.698 µg/g chlorophyll a, 0.263 µg/g chlorophyll b, 0.147 µg/g carotene, 1.620 µg/g total carotenoid and 1.473 µg/g xanthophyll contents in treated samples was statistically significantly different from control value of 4.366 µg/g chlorophyll a, 3.132 µg/g chlorophyll b, 0.796 µg/g carotene, 17.755 µg/g total carotenoid and 16.937 µg/g xanthophyll contents by day 50 samples. After prolonged exposure, growth rate, yield and pigment content for the treated samples recovered to control levels on day 120. The obtained data indicate the application of aquatic plant *Lemna gibba* as sensitive biomarker of water quality as well as the significance of selected biological parameters in the reliable assessment of toxic potential of MSM under rice cropping condition.

## 1. Introduction

Metsulfuron methyl a sulfonylurea herbicide has very low mammalian toxicity and is widely used in agriculture (James 1990). MSM has a potential to contaminate the ground water at very low concentration (Petkova and Donkova 2003) and at the same time, aquatic species are greatly affected by the presence of pollutants in surface water (Roy and Hanninen 1994; Pflugmacher et al. 1999; Hand et al. 2001). MSM was environmentally companionable and require very small quantity to kill the weeds by inhibiting the enzyme acetolactate synthase(ALS), which is necessary in the first step for plants to synthesize the branched amino acids, valine, leucine and isoleucine (Brown 1990). The degradation reaction of MSM in the surface of plant directly impacts the herbicide effectiveness (Eyheraguibel et al. 2011). Hence, the potential impact of contaminants on primary producers and their environmental risks is assessed using aquatic plant toxicity test (Fairchild et al. 1997). The *Lemna gibba* is commonly used in phytotoxicity tests because of its small size, high reproductive rate, ease of cultivation and ease of growth measurement without specified instruments (Wang 1990). The growth and reproduction of *Lemna* species is faster than other vascular plants in the aquatic phytotoxicity assessment tests (EPA 1996; OECD 221 2002). Sulfonylurea herbicide (SUH) inhibits rapid cessation of plant cell division and growth. SUH and its metabolite retaining the sulfonylurea function were highly toxic to *Lemna gibba*, whereas a metabolite lacking the sulfonylurea function was practically non-toxic (Marks 2008). The pigment content in *Lemna* species is one of the first visible symptoms. These changes are used as an indicator of photosynthetic damage in plant tissues. Biomarkers such as pigment content (chlorophyll, carotenoid, carotene and xanthophylls) are commonly used as parameters for toxicity tests (Mohan and Hosetti 1999). The pigment content in *Lemna* species is to identify the toxicity of SUH (Dos Santos et al. 1999). Hence, the present study the impact of MSM and its residues on the growth of *Lemna* has been investigated. The main objectives of the present study were to assess the impact of residues on the growth and yield of *Lemna gibba* and its recovery at different occasions under actual rice cropping condition, to determine the potential toxicity Index value of MSM to *Lemna gibba* and to investigate the effect of MSM on pigment content in the *Lemna* species.

## 2. Materials and Methods

### 2.1. Sources of Chemicals and Test System

Reference standard of MSM (purity 98%) was purchased from Sigma Aldrich, USA. MSM formulation (20% WG - water dispersible granule) was purchased from local vendors. All other chemicals are HPLC grade chemicals obtained from Merck India Limited, unless specified. *Lemna gibba* culture, maintained in the Department of Ecotoxicology, International Institute of Biotechnology and Toxicology, Padappai, Tamil Nadu, India was used in the study.

### 2.2. Experimental

#### 2.2.1. Residue analysis

The residues of MSM in water sample was studied by applying the herbicide formulation at a recommended dose of 8 g a.i /ha in duplicate. Rice seeds (ADT-43) were sown in a plot (10 sq. m.) located at International institute of biotechnology and toxicology (IIBAT), Padappai near Chennai. The age of rice used was 10 days crop after transplanting at the time of spray. The required water level (4 — 5 cm) was maintained throughout by frequently irrigating the field upto 120 days. On each sampling occasion (0, 5, 10, 15, 30, 50, 70, 100 and 120 days), water samples were collected from the field, filtered using a 0.2µm polytetrafluoroethylene (PTFE) membrane filter and analyzed for residue and its metabolite analysis by using validated HPLC-UV and Liquid Chromatography Electrospray Tandem Mass Spectrometry (LC-MS/MS-ESI) method.

#### 2.2.2. *Lemna* assay test

##### 2.2.2.1. Pre-culture

Young, rapidly growing *Lemna* plants was used in the study. Sufficient colonies of *Lemna gibba* culture was transferred aseptically into fresh sterile culture medium (20X AAP medium (OECD 221)) for 7 days before initiation of the study. The inoculated flasks were kept in the growth cabinet as slanting position providing continuous illumination of 6888–7132 lux intensity using white fluorescent lamps with temperature ranging from 22.4–23.1°C for 7 days. *Lemna gibba* were pre-cultured in fresh sterile culture for 7 days before the actual exposure to MSM-treated water under rice cropping condition.

##### 2.2.2.2. Test Procedure

An aliquot of 160 ml water sample was collected in a 500 ml beaker from exposed samples in rice-soil-water ecosystems occasionally. The control and treated samples were inoculated with equal number of *Lemna gibba* plants (10 fronds) from a 7 days old pre-culture under aseptic conditions in triplicate. The test was terminated 7 days after transferring the plants into the test vessels. During the experimental period, the temperature in the test medium was 27.3–41.2°C and the light intensity was recorded with the range of 45000 — 60000 lux under direct sunlight. Number of *Lemna* fronds was recorded for both treated and control at pre-determined intervals and calculated the percentage of inhibition of yield. Every frond visibly projecting beyond the edge of the parent frond was counted. Observations on the appearance of the fronds included changes in plant size or shape and root growth. The morphological changes were further confirmed by the estimation of chlorophyll a, chlorophyll b, carotenoid, carotene and xanthophylls content in *Lemna* species using UV-Visible spectrophotometer method.

##### 2.2.2.3. Evaluation of Percentage of Inhibition on growth rate and Yield of Lemna gibba

The effect of MSM-treated water from field condition on the vegetative growth of *Lemna* was evaluated based on assessments of frond number. To quantify the substance related effects, growth in the test solutions were compared with control and the concentrations brought about a specified percentage inhibition of yield based on frond numbers. The percentage inhibition of average specific growth rate and yield (%*I*_*r*_ and % *I*_*y*_) was calculated as, 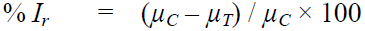 where, %*I*_*r*_ = percent inhibition in average specific growth rate

*µ*_*C*_ = mean value for *µ* (average specific growth rate) in the control

*µ*_*T*_ = mean value for *µ* (average specific growth rate) in the treatment group

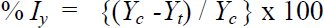

Where *I*_*y*_ - percent inhibition of yield based on frond numbers

*Y*_*c*_ - mean value for yield in the control

*Y*_*t*_ - mean value for yield in the treatment group

The doubling time of frond number in the control was 2.03 days which is less than 2.5 days (60 hours) and the pH of the control field is 8.38 which was recorded in the limit of 1.5 units during the test. Based on the results, the study is validated.

#### 2.2.3. Estimation of Toxicity Index *(TI)*

TI values were evaluated as the concentration of MSM in the sample at pre-determined intervals divided by the EC_50_ or LC_50_ (half of the effective concentration/lethal concentration) of an aquatic organism. Acute toxicity data of sulfonylurea herbicides were primarily collected from the EPA’s ECOTOX database (EPA, 2001) and other sources (Fairchild et al., 1997; Sparling et al., 2000). Published EC_50_ value of MSM (0.38 µg/L) for aquatic plant *Lemna gibba* was used as toxicity metrics in this present study (Battaglin and Fairchild, 2002). The Toxicity Index is calculated as follows: TI value = Concentration of MSM in different occasional samples / EC_50_value of MSM; If, TI values > 1.0 indicates probable toxicity to aquatic plant; > 0.5 indicates potential toxicity to aquatic plant and > 0.1 indicates limited toxicity respectively.

#### 2.2.4. Analysis of Pigment content

Approximately 200 mg of *Lemna* species was taken in a mortar and pestle and homogenized with 10 ml of 95% methanol. The homogenized sample was filtered through a Whatman filter paper 41 and the absorbance of the filtered solution was measured at 666, 653, 470 and 436 nm (UV-Visible Spectrophotometer, Shimadzu) for chlorophyll a, chlorophyll b, total carotenoid and carotene content. The quantities of pigments (chlorophyll a, chlorophyll b, total carotenoid, carotene and xanthophylls) were calculated according to the given formula with different solvents such as acetone, methanol and diethylether (Lichtenthaler and Wellburn, 1985). Data was compared by analysis of Student’s t-test at alpha level 0.05 using SAS 9.3 software was performed to determine the significant difference between treatment and control. Each data point is the average of three replicates (n=3), unless stated otherwise.

## 3. Results and Discussion

### 3.1. Residue analysis

The HPLC method for the determination of residue concentration had an acceptable recovery 70-120% in water sample by fortifying two different concentrations of MSM at 100 and 10 µg/L. The limit of detection (LOD) and limit of quantification (LOQ) of MSM established were 10 µg/L.The MSM residue in field water was determined on day 0, 5, 10, and 15 were 1254, 756, 125 and 11µg/L, respectively. The presence of MSM residues by day 30 was below detectable level (BDL). The details of MSM and metabolite / breakdown products are presented in **Table 1**. The use of analytical technique for residue concentration was very difficult at below the quantification level. Hence, the potential impact of MSM residue after day 15 was further confirmed by using growth inhibition test. The aquatic plant *Lemna gibba* was used to assess the impact of MSM residue at predetermined intervals under actual cropping condition.

**Table 1.** LC-ESI-MS/MS fragmentation ions of metsulfuron methyl and its breakdown products

| Compound name | Molecular mass | Positive Mode |  | Negative Mode |  |
| --- | --- | --- | --- | --- | --- |
|  |  | Molecular Ion | Fragment ion | Molecular Ion | Fragment ions |
| Metsulfuron methyl | 381 | 380 | 139, 182, 214 | - | - |
| DesmethylMetsulfuron | 367 | 368 | 127, 153, 199 | 366 | 125, 151, 182 |
| Methyl-2-(carbamoylsulfamoyl)benzoate | 258 | - | - | 257 | 182, 214 |
| 4-methoxy-6-methyl-1,3,5-triazin-2-amine | 140 | 141 | 58, 85, 124 | - | - |
| 4-amino-6-methyl-1,3,5-triazin-2-ol | 126 | 127 | 86, 60 | - | - |

### 3.2. Estimation of percentage Inhibition of growth and yield on *Lemna gibba*

During the experimental period, the temperature in the test medium was 27.3 — 41.2°C and the light intensity was recorded with the range of 45000 — 60000 lux under direct sunlight. MSM caused maximum inhibition (100%) on the growth and yield of *Lemna* from day 0 to day 50. The above inhibition was due to the presence of parent compound and its breakdown products retaining sulfonylurea group. In the literature cited, MSM and its metabolite retaining the sulfonylurea function were highly toxic to *Lemna gibba*. MSM was inhibiting ALS enzyme to stop the synthesis of branched amino acids, valine, leucine and isoleucine for the first step of plant growth, root elongation and finally plant was dead. The above symptom was observed during the period between 0 and 50 days. On the 70^th^ day, the percent inhibition on growth rate and yield was 73% and 78%, respectively. The lowest value of inhibition on growth and yield on the 100^th^ day was 22% and 26%. The above data indicate the degradation of MSM and breakdown products retaining sulfonyl urea group and *Lemna* growth was recovered slowly. Analysis of the 120^th^ day samples showed no sign of inhibition on the growth and yield of *Lemna gibba*, which is recovered to control levels. Results of the study revealed that, sulfonylurea herbicide MSM at recommended dose shows substantial inhibitory effect when compared to control on the growth rate and yield of *Lemna gibba*.

### 3.3. Estimation of TI value

In the present study, the additional parameter as TI value was calculated to measure the toxic effect of MSM residue. TI value is greater than 1.0 till the 15^th^ day, which indicates that the concentration of MSM in water sample was highly toxic to *Lemna gibba* due to parent compound MSM and its metabolites having sulfonylurea group. The residual data of MSM in water sample under actual cropping condition, percentage inhibition of growth and yield based on frond numbers of *Lemna gibba* and toxicity index value was calculated during the period on different occasions are presented in **Table 2**.

**Table 2.**
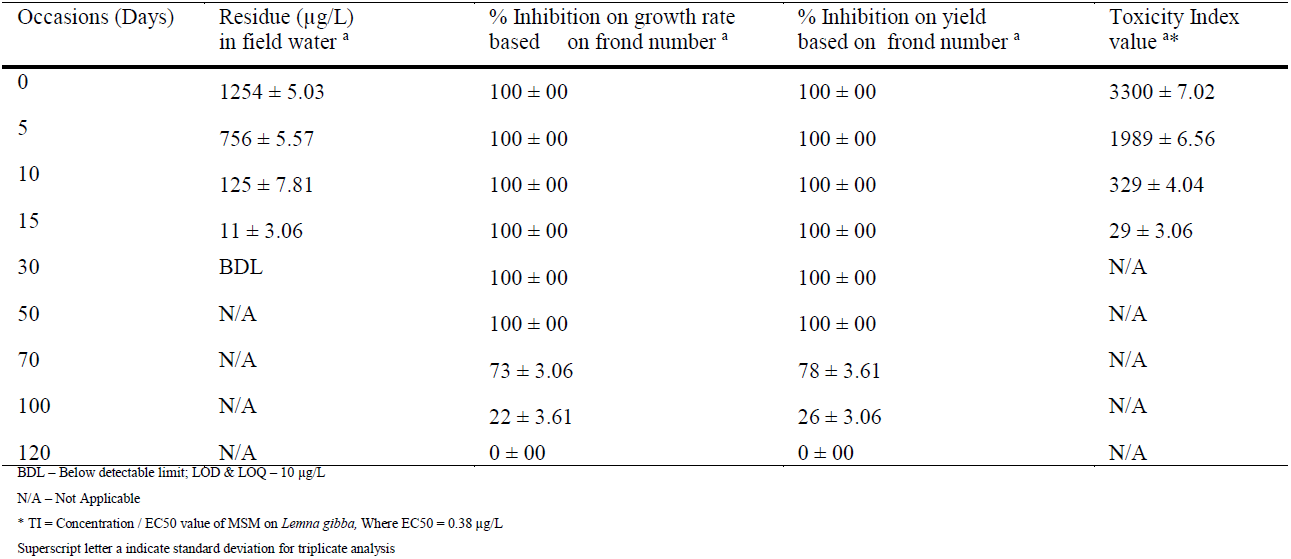
Correlation of residue concentrations of MSM under cropping condition with percentage inhibition on growth rate and yield and toxicity index value for over a period of four months

| Occasions (Days) | Residue ($\mu\text{g/L}$)<br>in field water <sup>a</sup> | % Inhibition on growth rate<br>based on frond number <sup>a</sup> | % Inhibition on yield<br>based on frond number <sup>a</sup> | Toxicity Index<br>value <sup>a*</sup> |
| --- | --- | --- | --- | --- |
| 0 | 1254 ± 5.03 | 100 ± 00 | 100 ± 00 | 3300 ± 7.02 |
| 5 | 756 ± 5.57 | 100 ± 00 | 100 ± 00 | 1989 ± 6.56 |
| 10 | 125 ± 7.81 | 100 ± 00 | 100 ± 00 | 329 ± 4.04 |
| 15 | 11 ± 3.06 | 100 ± 00 | 100 ± 00 | 29 ± 3.06 |
| 30 | BDL | 100 ± 00 | 100 ± 00 | N/A |
| 50 | N/A | 100 ± 00 | 100 ± 00 | N/A |
| 70 | N/A | 73 ± 3.06 | 78 ± 3.61 | N/A |
| 100 | N/A | 22 ± 3.61 | 26 ± 3.06 | N/A |
| 120 | N/A | 0 ± 00 | 0 ± 00 | N/A |

BDL – Below detectable limit; LOD & LOQ – 10 $\mu\text{g/L}$
N/A – Not Applicable
\* TI = Concentration / EC50 value of MSM on *Lemna gibba*, Where EC50 = 0.38 $\mu\text{g/L}$
Superscript letter a indicate standard deviation for triplicate analysis

### 3.4. Photosynthetic pigments

The changes in pigment content on *Lemna* plant is one of the first visible symptoms. These changes are used as an indicator of photosynthetic damage in plant tissues. The above morphological changes were confirmed by the analysis of pigment content in the plant using spectrophotometer. Pigment content of the control plants ranged from 4.319-4.371 µg/g (Chlorophyll a), 3.106-3.162 µg/g (Chlorophyll b), 0.782-0.811 µg/g (carotene), 17.653-17.758 µg/g (carotenoid) and 16.904-16.948 µg/g (xanthophylls) throughout the study was presented in **Fig. 1A**. The total pigment content in *Lemna* was affected on the 0 day sample, after 7 days exposure of *Lemna gibba*. Significant decrease of pigment content was observed during period between day 0 and day 50. The value of pigment content on Day 50 was 0.698 µg/g chlorophyll a, 0.263 µg/g chlorophyll b, 1.620 µg/g carotenoid, 0.147 µg/g carotene and 1.473 µg/g xanthophyll content, which indicates the destruction of photosynthetic pigments by MSM residue due to deficiency of electron transport chain, replacement of Mg^2+^ ions associated with the tetrapyrrole ring of chlorophyll molecules by H^+^ ion, inhibition of important enzymes (Van Assche and Clijsters 1990) for the synthesis of chlorophyll molecules or peroxidation processes in chloroplast membrane lipids by the reactive oxygen species (Sandalio et al. 2001). On the 120^th^ day, chlorophyll a, chlorophyll b, carotene, carotenoid and xanthophyll content were on par with control, as is shown in **Fig. 1B**. The total pigment values on day 120 indicated that L*emna* had recovered the growth.

**Fig. 1.**
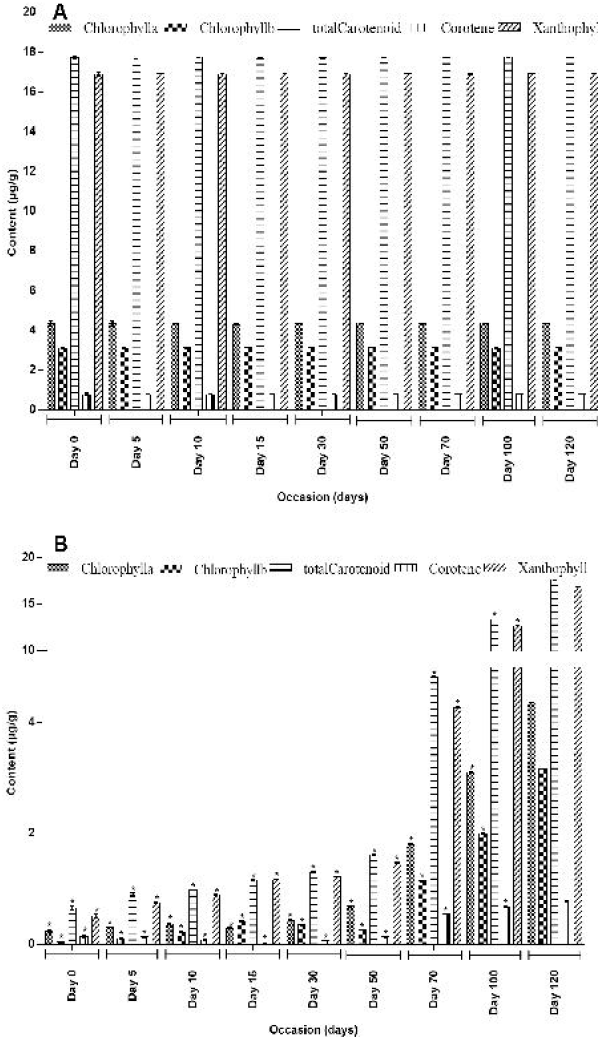
A represents pigment contents in control samples and B represents pigment contents in MSM treated samples. Relative chlorophyll *a*, chlorophyll *b*, carotene, total carotenoid and xanthophyll contents in aquatic plant *Lemna gibba* exposed to 7 days to field water samples collected pre-determined intervals over a four months period. Standard deviations were presented by *error bars*. Values are mean of three replicates. Symbol * indicate significantly different from control upto 100 days (α<0.05) and recovered on day 120.

## 4. Conclusion

The results show the suitability of *Lemna gibba* for field water quality assessment as all selected parameters showed impact of MSM residue with respect to water samples collected with different occasions over a four month period. The result is large variability between control and treated individuals. This variability is attributed due to the presence of trace level of residues of MSM and its breakdown products retaining sulfonylurea group in water samples. The decline in total chlorophyll, carotene, carotenoid and xanthophyll contents as well as growth inhibition can be regarded as general response associated with MSM toxicity. After prolonged exposure by day 120, growth rate, yield and pigment content of *Lemna* was recovered, which is on par with control. The present assessment also highlights the successful application of *Lemna gibba* as a potential plant biomarker to identify the impact of herbicide residues at trace level. The outcomes attained suggest that phytotoxicity tests with aquatic plant *Lemna gibba* should be used in the biomonitoring of field water due to its simplicity, sensitivity and cost-effectiveness.

## Acknowledgement

The authors are thankful to the Management, International Institute of Biotechnology and Toxicology for providing the necessary facilities to perform this research work.

